# Upcycling Polyethylene into Poly(3-hydroxybutyrate) via a Chemo-Enzymatic-Microbial Cascade

**DOI:** 10.1101/2024.09.26.615289

**Authors:** Demin Kong, Lei Wang, Wei Xia, Meng Shi, Qisheng Fu, Guiyuan Zheng, Jing Wu

## Abstract

Polyethylene (PE) plastics are extensively utilized across agricultural, industrial, and medical sectors owing to their favorable physicochemical properties. However, their chemical stability and escalating production have resulted in severe waste accumulation and environmental pollution. Conventional disposal methods are plagued by resource inefficiency and secondary pollution. While emerging strategies offer promise, physicochemical methods demand harsh operating conditions, and biological routes remain inefficient. This research presents an integrated “chemical pretreatment–biodegradation–upcycling” system that combines the efficiency of chemical catalysis with the sustainability of biological conversion. Ester bonds were introduced into PE via Baeyer–Villiger oxidation, followed by enzymatic hydrolysis using the cutinase from *Thermobifida fusca* WSH03-11 (*Tf*Cut). Specifically, machine learning-aided optimization of reaction conditions and computational redesign of *Tf*Cut enhanced degradation efficiency, yielding a maximum weight loss of approximately 71.19%. The degradation intermediates were bio-converted into poly(3-hydroxybutyrate) (PHB) by the wild-type strain LETBE-HOU, isolated in this study, achieving a concentration of 16.75 mg/L. Multi-omics analysis of LETBE-HOU further revealed the PHB biosynthesis pathway and fatty acid degradation regulation. This work breaks the long-standing reliance on physiochemically-derived degradation intermediates for microbial conversion of PE, establishing a fully circular system that opens new avenues for future research.

## Introduction

Polyethylene (PE), the most prevalent member of the polyolefin family, is a high-molecular-weight polymer composed of ethylene monomers connected by carbon-carbon single bonds. Its attractive properties, including abundant availability, low production cost, and favorable physicochemical characteristics, have driven continuous global production expansion. Cumulative output between 1950 and 2017 surpassed 180 million tons, with projections indicating an annual growth rate of 2-5%, potentially exceeding 2.5 billion tons by 2050^1, 2, 3^. This escalating production has generated correspondingly large amounts of plastic waste. However, the carbon-carbon bonds in PE exhibit high dissociation energy (about 330-370 kJ/mol), substantially greater than that of natural polymers such as lignin or latex^4, 5^. This exceptional stability renders PE highly persistent in the environment, with estimated degradation timescales of up to 400 years^6^. The convergence of chemical recalcitrance and unabated production growth has intensified waste accumulation and environmental pollution, underscoring the critical need for efficient and sustainable degradation strategies, particularly through advanced biological and catalytic conversion pathways.

Current disposal methods for PE predominantly rely on incineration and landfilling, which approaches that present considerable environmental drawbacks. Incineration generates significant secondary pollutants, while landfilling consumes vast tracts of land, requires extended mineralization timelines, and facilitates the continuous release of microplastics. Alternative physicochemical techniques, including pulverization, decrystallization^7^, ultraviolet irradiation^8^, catalytic cracking^9, 10, 11^, and microwave-assisted catalysis^9, 10, 12, 13^, can decompose PE into alkanes, alkenes, and other chemical products. Despite their high catalytic efficiency, these methods are constrained by operational safety issues and demanding reaction conditions, hindering large-scale implementation. Biological degradation methods, mediated by whole cells^5^ or isolated enzymes^14^, provide compelling advantages such as mild operating conditions and inherent environmental compatibility. However, they face a fundamental limitation: the recalcitrance of carbon-carbon backbone to direct enzymatic cleavage. Drawing inspiration from physicochemical processes, where intensive oxidative treatments introduce functional groups and reactive sites into the polymer matrix, a synergistic strategy has emerged^15^. By combining initial physicochemical pretreatment with subsequent biological conversion, it becomes possible to enhance bioavailability and degradation efficiency of PE^8, 16, 17, 18, 19, 20^. This integrated approach represents a promising and sustainable platform for plastic valorization, aligning degradation performance with green process ideals.

Pretreatment of PE centers on the introduction of polar functional groups into its polymer backbone to enhance degradability. These reactive sites encompass a range of polar moieties, such as hydroxyl, carbonyl, C=C double bonds, and ester bonds^21^. Among them, ester bonds represent a particularly viable target owing to their well characterized enzymatic cleavage and higher practical feasibility^21^. Following established approaches^22, 23^, ester groups can be incorporated into PE via Baeyer–Villiger (BV) oxidation, and subsequent enzymatic hydrolysis by the cutinase from *Thermobifida fusca* WSH03-11 (*Tf*Cut) enables effective depolymerization. Nevertheless, prior studies have not accomplished closed–loop recycling of the resulting degradation intermediates.

Specifically, current research on the valorization of PE degradation products focuses on three main routes: mineralization^6^, conversion into small molecules^20^, and biosynthesis of biodegradable polymers^16, 24^. The microbial conversion of PE degradation intermediates into low-molecular-weight compounds or biodegradable plastics represents a promising route toward efficient and sustainable plastic waste management. However, in most reported cases^6, 16, 24^, the substrates metabolized by microorganisms are primarily derived from physicochemical degradation processes. To date, the direct reuse of PE biodegradation intermediates remains unreported, underscoring a critical research gap and highlighting the need for integrated processing strategies.

To tackle the challenges outlined above, this work develops an integrated “degradation–upcycling” platform (Fig. 1) that enables the complete biological conversion of PE into the biodegradable plastic poly(3-hydroxybutyrate) (PHB). System performance was substantially enhanced through machine learning-assisted optimization of reaction parameters and computational redesign of the key enzyme *Tf*Cut, yielding a notable weight loss of up to 71.19%. By employing PE degradation intermediates as the sole carbon source, we isolated a novel wild-type strain, LETBE-HOU, capable of synthesizing PHB at titers reaching 16.75 mg/L. The degradation efficiency achieved here exceeds current benchmarks for biological PE degradation systems^19, 25, 26, 27, 28^. More significantly, this study achieved the first microbial valorization of high-molecular-weight PE degradation intermediates (>2000 Da), thereby overcoming a fundamental limitation in a field where only low-molecular-weight fragments from physicochemical pretreatment were previously considered metabolizable^6, 16, 24^. Combined with systematic pathway analysis of PHB synthesis in LETBE-HOU, our findings provide both mechanistic insights and an engineered platform for sustainable plastic upcycling.

**Fig. 1.**
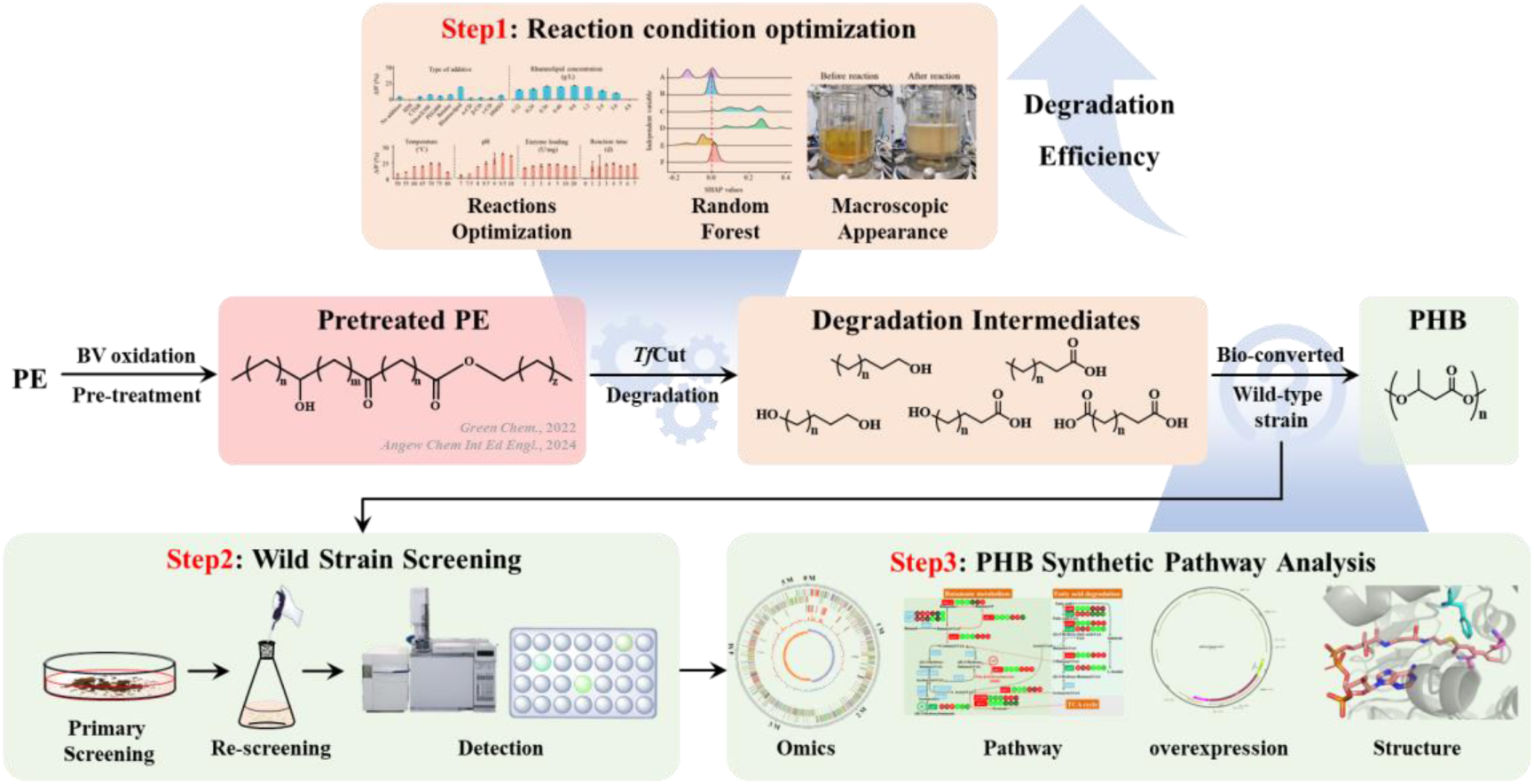
Schematic overview of PE waste upcycling via ester bond activation. The upcycling process comprises four stages: (1) Optimization of TfCut reaction conditions; (2) Screening of wild-type strains using PE degradation intermediates as the sole carbon source; (3) Functional characterization of the selected strain. Stages 1 and 2 aim to maximize PE degradation efficiency, while stages 3 and 4 enable upcycling of degradation intermediates into value-added products.

## Result

### Pathway for PE conversion to PHB

Prior studies have established that PE can be degraded into fatty acids, fatty alcohols, α,ω-carboxylic acids, and ω-hydroxycarboxylic acids via BV oxidation and lipase-catalyzed hydrolysis^22, 23^. These degradation intermediates serve as precursors for PHB synthesis^16^. Building on this foundation, an integrated reaction pathway was designed for converting PE to PHB, as illustrated in Fig. 1. Previous attempts in this area have encountered limitations such as marginal weight loss and inability to valorize degradation intermediates. To address these challenges, reaction parameters were optimized using random forest modeling, and novel microbial strains capable of transforming PE degradation intermediates into PHB were isolated. Collectively, this work establishes a comprehensive and efficient PE recycling system, successfully demonstrating a bio-based strategy for PE biodegradation and upcycling.

### Pre-treatment and degradation of PE via chemo-enzymatic cascade

The BV oxidation of PE was catalyzed by *^m^*CPBA to introduce ester bonds into the polymer backbone^22^, converting recalcitrant carbon-carbon single bonds into more labile ester linkages. Following established procedures^22, 23^, PE was pretreated with *^m^*CPBA and subsequently degraded using *Tf*Cut. The molecular weight profile and composition of the resulting degradation products aligned with previous reports^22, 23^. Under baseline conditions, *<u>Tf</u>*Cut-mediated degradation achieved a weight loss (Δ*W*) of approximately 4.34% (Fig. 2a), indicating limited degradation efficiency. However, the low extent of degradation hindered effective recovery and upcycling of the intermediates, necessitating strategies to enhance the overall degradation yield.

**Fig. 2.**
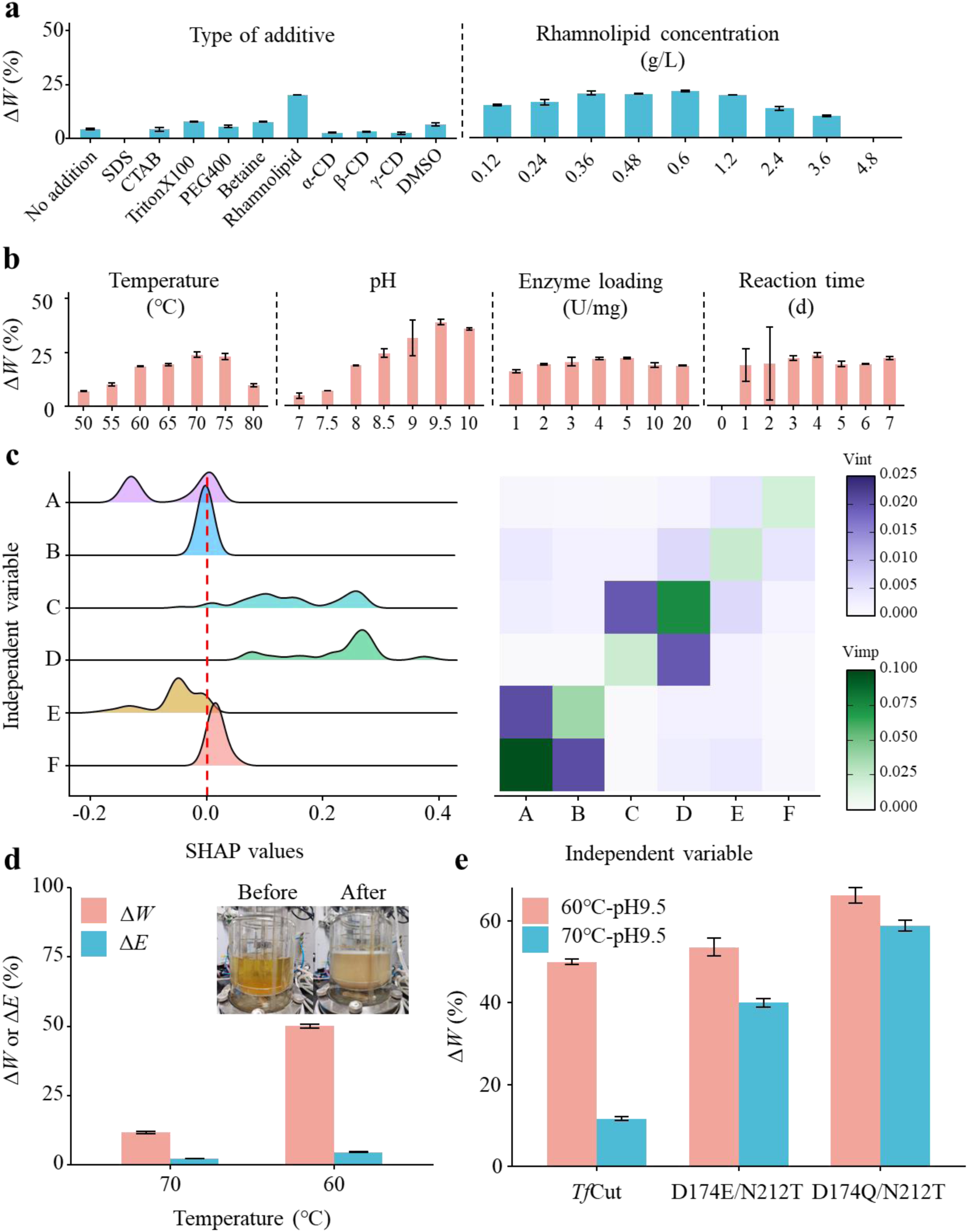
Optimization of reaction conditions of *Tf*Cut degradation system. **a,** Single-factor optimization of additive type and concentration. **b,** Single-factor optimization of temperature, pH, enzyme loading, and reaction time under fixed rhamnolipid concentration (0.60 g/L). **c,** Analysis of random forest modeling for single-factor optimization results via SHAP, vimp, and vint. Independent variables A–F correspond to reaction time, enzyme loading, pH, temperature, rhamnolipid concentration, and additive type, respectively. **d,** Degradation efficiency after multi-factor combinations (left) and macroscopic appearance of the reaction system before and after degradation (right). **e,** Δ*W* of pretreated polyethylene by wild-type *Tf*Cut and selected mutants. In the figure, Δ*W* represents the weight loss rate, and Δ*E* represents the enzyme activity residual rate, with calculation methods detailed in the Supplementary Methods 7.

### Reaction condition optimization to enhance degradation efficiency

Rational design of enzymatic catalytic systems has been shown to substantially improve the degradation efficiency of polymeric substrates^29, 30^. In this study, the *Tf*Cut-based degradation system was optimized through systematic refinement of key parameters including additive type and concentration, rotational speed, enzyme loading, temperature, pH, and reaction duration. Single-factor optimization (Fig. 2a,b) identified the following optimal conditions: 0.60 g/L rhamnolipid (22.04% weight loss), 70°C (35.37%), pH 9.5 (38.50%), enzyme loading of 5 U/mg substrate (22.14%), and 1-day reaction time (18.96%).

Random forest modeling was subsequently employed to evaluate the relative importance of multiple reaction parameters on degradation efficiency, providing a data-driven foundation for multi-factor optimization. Following dataset preprocessing and parameter tuning (Fig. S1), the model demonstrated strong predictive performance with correlation coefficients of 0.930 (training set) and 0.938 (test set) between predicted and actual values (Fig. S2). SHapley Additive exPlanations analyses (SHAP), variable importance (vimp), variable interaction (vint), partial dependence plot (PDP), and individual conditional expectation (ICE) (Fig. 2c and S3) consistently identified pH as the most significant factor, guiding its prioritization in subsequent experimental designs. The final optimized system, comprising 0.60 g/L rhamnolipid, 60°C, pH 9.5, 5 U/mg enzyme loading, and 1-day reaction, achieved a maximum weight loss of approximately 50.04% (Fig. 2d). Macroscopic examination revealed an emulsion-like state in the post-reaction mixture, indicating substantially improved aqueous solubility of the degradation products (Fig. 2d).

Comprehensive characterization of the degradation products was performed using differential scanning calorimetry (DSC), Fourier transform infrared spectroscopy (FT-IR), high-temperature gel permeation chromatography (HT-GPC), and gas chromatography–mass spectrometry (GC–MS) (Fig. S4). Compared with previous reports^22, 23^, the products exhibited reduced ester bond content, increased hydroxyl group abundance, lower molecular weight, higher ester bond utilization efficiency, and greater diversity of liquid-phase compounds. These results indicated that the optimized *Tf*Cut catalytic system can degrade PE more efficiently, offering greater potential for recycling.

However, system optimization revealed operational limitations, particularly rapid inactivation of *Tf*Cut under extreme alkaline conditions (pH 9.5) (Fig. 2ab and S5). To address this constraint, protein engineering was undertaken to enhance *Tf*Cut stability under high-temperature and high-pH conditions. Following previous studies, this study further employed the stability-enhanced mutant D174Q/N212T for the degradation of pretreated PE. Degradation performance showed that both mutants exhibited superior activity under pH 9.5/60 °C and pH 9.5/70 °C, with D174Q/N212T achieving the highest weight loss rate of ∼66.32% (Fig. 2e). Furthermore, through more precise temperature and pH control, the maximum weight loss rate in a 1 L reactor was increased to 71.19%.

### Screening and assessment of wild-type strains for valorization PE degradation intermediates

Wild-type strains were screened for their capacity to utilize PE degradation intermediates as carbon sources. Primary screening employed solid plates supplemented with Nile red to identify isolates with metabolic activity toward the intermediates. From this process, 15 wild strains were isolated, with representative plate cultures shown in Fig. S6. Subsequent liquid culture assays measured OD_600_, reflecting growth, and Nile red fluorescence intensity, indicating accumulation of polyhydroxyalkanoate-like compounds. As summarized in Fig. 3a,b, nine strains exhibited robust growth, and one isolate (ID 1) showed fluorescence intensity more than twice that of others. The high-performing strain, designated LETBE-HOU, demonstrated superior potential for polyhydroxyalkanoate synthesis and was selected for further analysis. Its growth curve was displayed in Fig. S7.

**Fig. 3.**
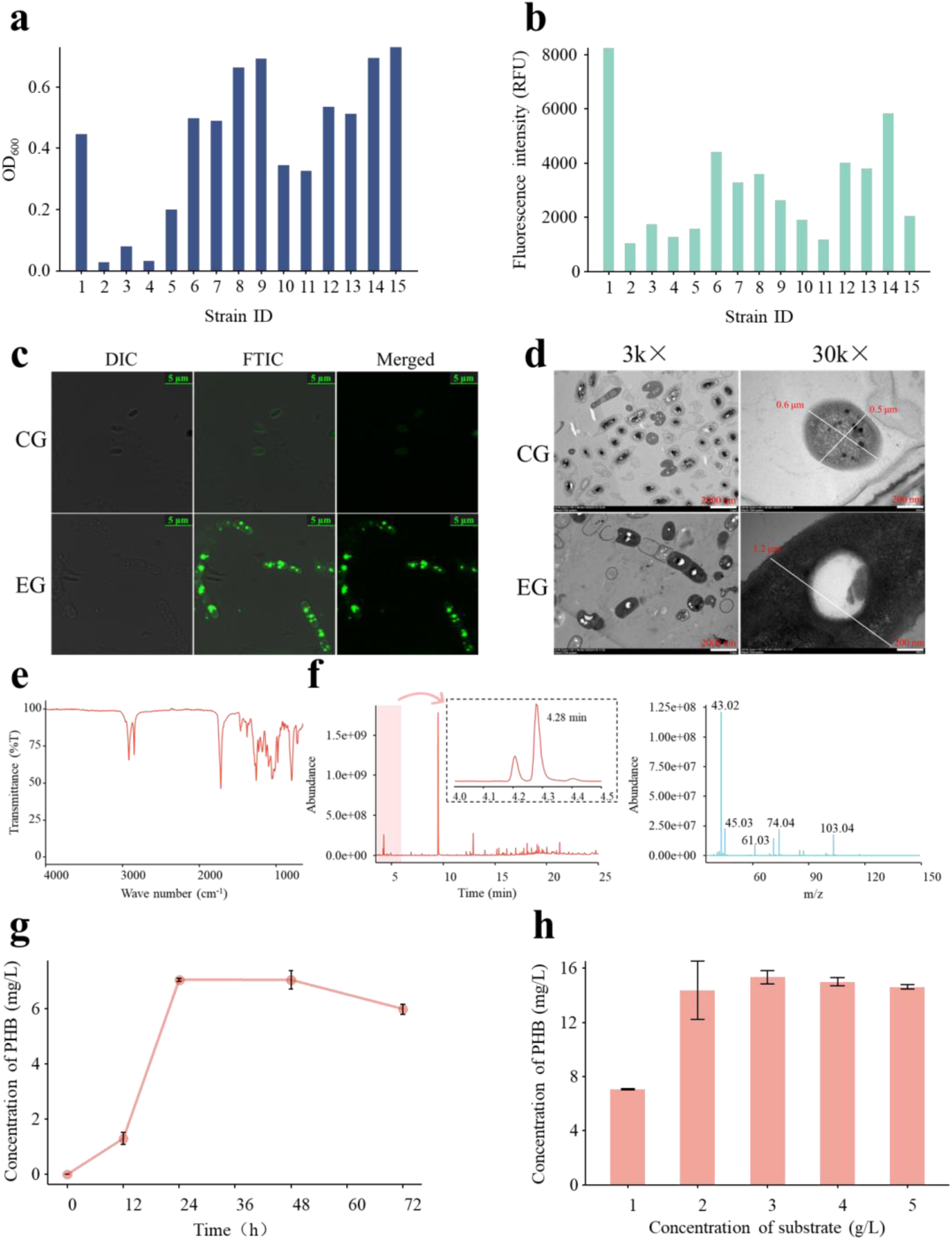
Screening of wild-type strains and characterization of metabolites from PE degradation intermediates. **a,** Growth status (OD_600_) of rescreened strains. **b,** Detection of metabolic product yield in strains via Nile red staining fluorescence. **c,** LSCM images of Nile red-stained LETBE-HOU cells. **d,** TEM micrographs of LETBE-HOU cells. In diagram b and c, EG and CG denote experimental and control groups, respectively. **e,** FT-IR spectrum of metabolites extracted from LETBE-HOU. **f,** GC–MS analysis of metabolites: chromatogram showing PHB monomer peak at 4.25 min and internal standard (methyl benzoate) at ∼9 min (left) with corresponding mass spectrum (right). **g,** Optimization of cultivation time for PHB production by LETBE-HOU. **h,** Optimization of PE degradation intermediate concentration for PHB synthesis.

Polymer production by LETBE-HOU was characterized using confocal laser scanning microscopy (LSCM), transmission electron microscopy (TEM), Fourier-transform infrared spectroscopy (FT-IR), and gas chromatography–mass spectrometry (GC–MS). LSCM (Fig. 3c) revealed bright intracellular fluorescent particles in cells cultivated with PE degradation intermediates (experimental group), indicating enhanced accumulation of lipid-soluble compounds relative to the control group. TEM (Fig. 3d) showed that experimental group cells were larger (1.2 μm) than control group cells (0.5 μm) and contained distinctly visible white granules within the cells, confirming intracellular metabolite accumulation. Microscopic examination revealed that this strain utilized PE degradation intermediates, with metabolic products accumulating intracellularly.

To chemically identify the accumulated product, FT-IR and GC–MS analyses were performed. The FT-IR spectrum (Fig. 3e) displayed a characteristic ester carbonyl absorption at 1730 cm^−1^, consistent with polyhydroxyalkanoates^31, 32^. GC–MS (Fig. 3f) confirmed the polymer as PHB. Under optimizing conditions (24 h cultivation, 3 g/L degradation intermediates), LETBE-HOU achieved a maximum PHB yield of 15.34 mg/L (Fig. 3g,h), exceeding previously reported values for similar strains^16, 24^. To our knowledge, this is the first reported strain capable of directly converting enzymatically derived PE degradation intermediates into PHB^16, 24^. The strain also efficiently assimilated long-chain fatty acids (Fig. S8), suggesting the presence of specialized transport or metabolic systems—a hypothesis meriting further genomic and transcriptomic investigation.

### Genome and transcriptome analysis for LETBE-HOU

To elucidate the PHB biosynthesis mechanism in strain LETBE-HOU, integrated genomic and transcriptomic analyses were conducted. Whole-genome sequencing identified the strain as *Bacillus paranthracis*, with an assembled genome has a total length of 5234090 bp and encodes 5198 genes. The circular genome map and a general summary are provided in Fig. S9 and Table S1, respectively. Functional annotation of the entire genome was subsequently performed using the Kyoto Encyclopedia of Genes and Genomes (KEGG), Gene Ontology (GO), Clusters of Orthologous Groups (COG), Carbohydrate-Active enZYmes Database (CAZy), and Pathogen Host Interactions Database (PHI) databases (Fig. 4a-e). KEGG annotation identified 167 membrane transport-related proteins; GO analysis showed enrichment in Catalytic Activity, Binding, and Transporter Activity within the Molecular Function category; and COG analysis highlighted numerous genes involved in Carbohydrate transport and metabolism and Cell wall/membrane/envelope biogenesis. These findings indicate a genetic predisposition for efficient transport and catalytic processes. CAZy annotation further identified genes encoding Carbohydrate-Binding Binding Modules (CBMs) and Carbohydrate Esterases (CEs), which may promote enzyme–substrate association and hydrolyze ester bonds in PE-derived intermediates, respectively. PHI annotation indicated that 13.70% of genes are linked to reduced virulence, supporting the strain’s suitability for biotechnological applications. Collectively, multi-database annotation underscores the genomic advantages of LETBE-HOU for PHB production, including efficient transport, catalytic, and binding systems, as well as robust cell envelope biogenesis.

**Fig. 4.**
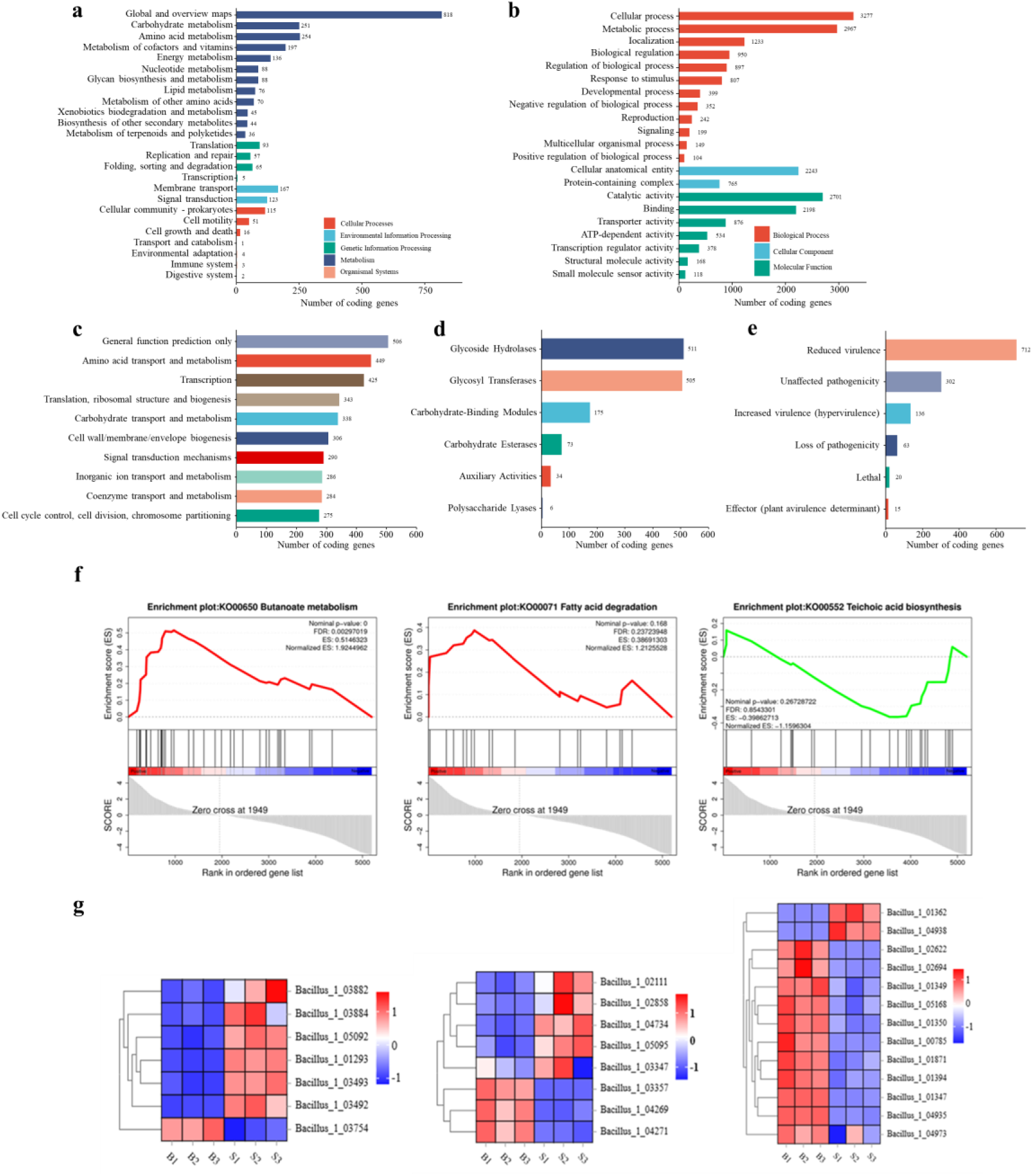
Genomic and transcriptomic analysis of LETBE-HOU. **a,** KEGG pathway annotation. **b,** GO classification. **c,** COG categorization. **d,** CAZy annotation. **e,** PHI analysis. **f,** GSEA of selected metabolic pathways: butyrate metabolism (left), fatty acid metabolism (middle), and peptidoglycan metabolism (right). **g,** Expression levels of key genes in butyrate metabolism (left), fatty acid metabolism (middle), and peptidoglycan metabolism (right) pathways. B and S denote the control and experimental group, respectively. All data were detected in triplicate.

Transcriptomic profiling under PHB-producing conditions revealed 930 upregulated and 1161 downregulated genes in the experimental group (Fig. S10). Gene set enrichment analysis (GSEA) of three PHB-relevant pathways—butyrate metabolism, fatty acid metabolism, and peptidoglycan metabolism—showed upregulation of the butyrate and fatty acid pathways and downregulation of peptidoglycan metabolism (Fig. 4f). Selected gene expression patterns from these pathways are displayed in Fig. 4g. Upregulation of the butyrate pathway, which is directly involved in PHB synthesis, and the fatty acid pathway, supplying key precursors, aligns with elevated PHB production. In contrast, downregulation of peptidoglycan metabolism likely reduces cell wall rigidity, potentially facilitating export of PE degradation intermediates and accommodating cell volume expansion during PHB accumulation. These transcriptomic results corroborate TEM observations and support a model in which LETBE-HOU reprograms its metabolism to efficiently convert PE degradation intermediate into PHB.

### Analysis and enhanced of PHB synthesis pathway for LETBE-HOU

Based on genomic and transcriptomic evidence, the PHB biosynthesis pathway in LETBE-HOU was elucidated. Key genes identified include those encoding NADPH-dependent acetoacetyl-CoA reductase (PhaB), 3-hydroxybutyrate polymerase (PhaC), poly-3-hydroxybutyrate inclusion body protein (PhaP), poly-3-hydroxybutyrate-responsive repressor (PhaQ), poly-3-hydroxybutyrate regulator (PhaR), and (R)-specific acyl-CoA hydratase (PhaJ). T Their genomic organization is consistent with previous reports^33^.Functionally, PhaP and PhaR regulate intracellular PHB stability, while PhaQ modulates PHB metabolic genes. The enzymes PhaB, PhaC, and PhaJ catalyze PHB biosynthesis.

PHB synthesis primarily occurs through the butyrate metabolism pathway (green region in Fig. 5a), via two distinct routes: 1) the conventional pathway: acetyl-CoA → acetoacetyl-CoA → (*R*)-3-hydroxybutyrate → PHB. Mainly catalyzed by PhaB and PhaC; 2) the auxiliary pathway: 2-butenyl-CoA → (*R*)-3-hydroxybutyrate → PHB, primarily involving the PhaJ and PhaC. Transcriptome analysis revealed that the most critical gene in this pathway is the PhaC-encoding gene (*phaC*), showing a 7.33-fold increase in expression compared to the control group (Fig. S11).

**Fig. 5.**
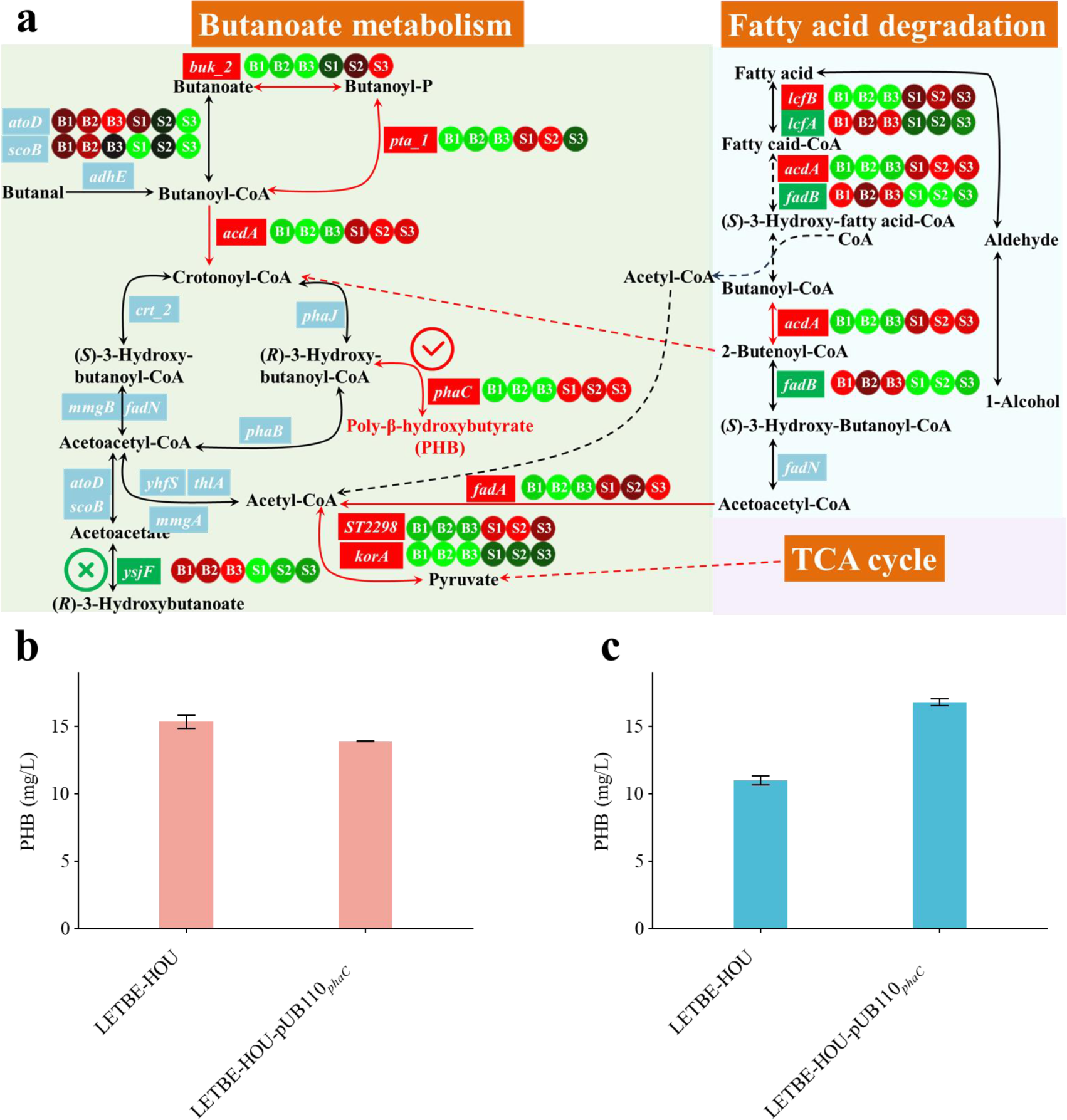
PHB metabolic pathway in LETBE-HOU and effect of phaC overexpression. **a,** PHB biosynthesis pathway from KEGG annotation: enzymes encoded by up-, down-, and non-regulated genes are highlighted in red, green, and blue, respectively; heatmap circles show expression levels (green to red: low to high). **b,** PHB production by *phaC*-overexpressing and wild-type strains using untreated PE degradation intermediates. **c,** PHB production by *phaC*-overexpressing and wild-type strains using decarboxylated PE degradation intermediates.

To validate its role, *phaC* was overexpressed in LETBE-HOU, and PHB production was evaluated using either untreated or redox enzyme CYP152N1-decarboxylated^34^ PE degradation intermediates. As showed in Fig. 5b,c, *phaC* overexpression did not enhance PHB yield with untreated substrates, likely due to insufficient raw material supply. However, with decarboxylated treatment, the engineered strain produced approximately 16.75 mg/L PHB, representing a 1.53-fold increase compared to the wild-type strain. These results confirm that *phaC* overexpression enhanced PHB production when precursor availability is sufficient. Further investigation into the substrate supply pathway (fatty acid metabolism) was required to improve the strain’s utilization efficiency of PE degradation intermediates.

### Analysis of fatty acid metabolism for LETBE-HOU

Further analysis of fatty acid metabolism revealed two major inconsistencies. First, a discrepancy was observed between GC–MS results and the theoretical fatty acid metabolic pathways. GC–MS analysis confirmed that LETBE-HOU exclusively produces PHB. In contrast, the canonical fatty acid metabolic pathway suggests that multiple acyl-CoA products can be converted by PhaJ to 3-hydroxyacyl-CoA, which PhaC then polymerizes into a mixture of polyhydroxyalkanoates (Fig. 3a). A second inconsistency was identified between the expression of a key fatty acid metabolism gene and observed bacterial growth. KEGG annotation identified only one acyl-CoA hydratase (FadB), whose expression was downregulated by 3.02-fold (Fig. S11). This downregulation insufficient to support the strain’s fatty acid metabolic demands in oligotrophic media.

Genomic and transcriptomic data mining identified an isoenzyme, acyl-CoA hydratase YhaR, which shares 27.45% sequence homology with FadB. AlphaFold3-based structural prediction and hexenyl-CoA docking revealed that that FadB and YhaR adopt conserved acyl-CoA hydratase folds but differ in active-site geometry (Fig. 6). In FadB, residue F141 creates a spatial barrier that confers specificity for short-chain acyl-CoAs (carbon chain length <6). In contrast, YhaR possesses an open conformation that accommodates long-chain acyl-CoAs (carbon chain length >6), with residue F137 at the substrate entrance acting as a gatekeeper to preclude short-chain substrates. Transcriptome data (Fig. S11) showed a 3.02-fold downregulation of FadB and a concurrent 7.56-fold upregulation of YhaR. These results support a “dual-enzyme control” model for fatty acid metabolism: YhaR regulates the stepwise conversion of long-chain fatty acids to 2-butenyl-CoA, while FadB precisely controls the degradation flux of 2-butenyl-CoA. This mechanism maintains overall fatty acid degradation rates while channeling 2-butenyl-CoA toward the specific synthesis of PHB.

**Fig. 6.**
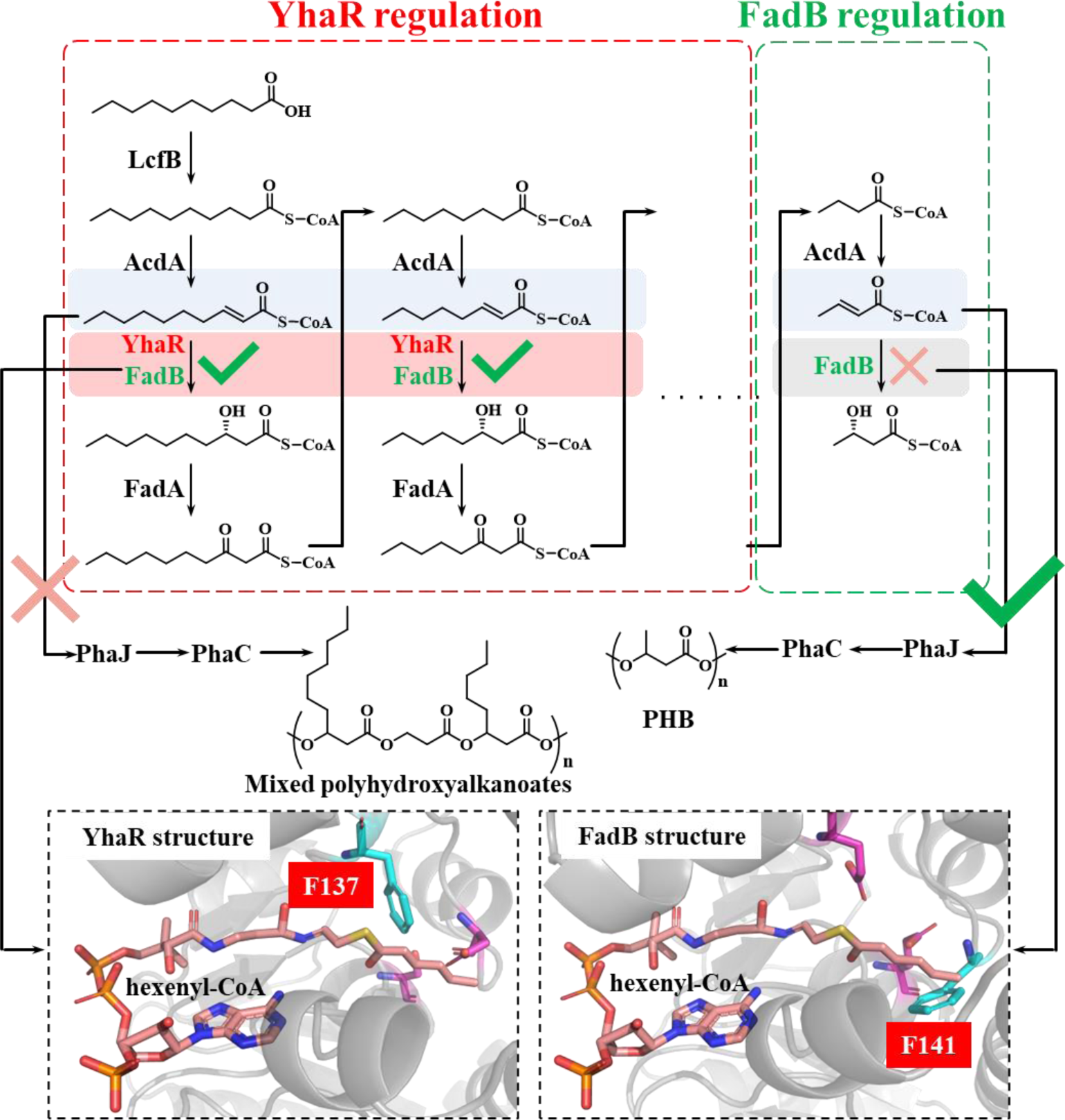
Schematic of the “dual-enzyme control” in fatty acid metabolism of LETBE-HOU.

This dual-enzyme control model provides a theoretical foundation for engineering the fatty acid metabolic. Nevertheless, further validation is required, including validation of the enzymatic functions of YhaR and FadB, investigation of the effects of gene knockout or overexpression on PHB production, and exploring the roles of additional acyl-CoA hydratase isoenzymes.

## Discussion

The persistent challenge of PE biodegradation necessitates innovative recycling strategies. While direct microbial or enzymatic degradation of PE remains elusive, this study establishes an integrated, three-stage recycling cascade comprising pretreatment (ester bond introduction), enzymatic degradation (ester bond hydrolysis), and microbial product conversion, thereby achieving the successful upcycling of PE. This system currently exhibits the highest catalytic efficiency reported for biocatalysis^19, 25, 26, 27, 28^. Simultaneously, each stage presents significant opportunities for optimization.

The current pretreatment system, which employs hazardous reagents and operates discontinuously from the subsequent degradation step, is a primary target for improvement. Substituting organic solvents with green alternatives like deep eutectic solvents (DES) and replacing *^m^*CPBA with enzyme-catalyzed (e.g., CALB) BV oxidation could mitigate environmental and safety concerns.

To address operational discontinuity, further engineering of cutinase—through immobilization, enhancing organic solvent tolerance, and improving stability—could enable a more seamless integration of the pretreatment and degradation stages into a unified, continuous process.

Regarding the PE degradation intermediates conversion stage, the strain LETBE-HOU requires further optimization to maximize PHB yield. Potential strategies include: engineering cellular morphology and membrane permeability to enhance intracellular PHB accumulation; strengthening substrate transport systems to improve uptake efficiency; expressing exogenous enzymes (e.g., oxidoreductases, esterases) to broaden substrate utilization; and modulating key PHB pathway genes to augment metabolic flux. These modifications are expected to significantly increase the overall conversion efficiency from PE to PHB. Notably, we observed that LETBE-HOU can survive as spores at 60°C, a trait that could be leveraged to facilitate its integration into a consolidated catalytic system encompassing both the pretreatment and degradation stages.

In summary, this work pioneers a novel chemo-enzymatic-microbial pathway for PE upcycling by converting it into the biodegradable polymer PHB, thereby laying a foundational framework for future research in plastic waste management.

## Methods

### Reagents, strain, and plasmid

Most Chemicals were procured from TCI, Sinopharm or Macklin. The bacterial strains and plasmids used in this study were obtained from previous studies^35^ or from our own screening. Detailed information is provided in Supplementary Methods 1.

### Enzyme preparation and characterization

The cutinase *Tf*Cut and its mutants were expressed in *Escherichia coli*. The recombinant proteins were purified via Ni-NTA affinity chromatography, and their stability and catalytic efficiency were characterized using model substrates. A detailed protocol is available in Supplementary Methods 2 and 3.

### PE Degradation system

The PE degradation process consists of two main stages. The pretreatment stage primarily employs *^m^*CPBA^22, 36^, to introduce oxygenated functional groups via simultaneous hydroxylation, carbonylation, and BV oxidation. The subsequent enzymatic degradation stage utilizes *Tf*Cut to hydrolyze the introduced ester bonds. The catalytic conditions for *Tf*Cut, including additive type/concentration, enzyme loading, temperature, pH, reaction duration, and stirring speed, were systematically optimized. Degradation efficiency was primarily quantified by measuring the weight loss of the residual solid material after the reaction. Further details are described in the Supplementary Methods 4 to 6.

### Detection of PE degradation products

The degradation products were characterized by DSC, FT-IR, HT-GPC, and GC–MS. These techniques were used to determine changes in melting temperature, functional groups, molecular weight, and degradation products, respectively. Detailed methodologies are described in the Supplementary Methods 7 to 11.

### Synthesis, Qualification, and Quantification of PHB

LETBE-HOU was obtained through a screening campaign of wild-type strains to synthesis PHB. It was pre-cultured in LB medium and subsequently transferred to YSV medium (containing 1 g/L PE degradation products) for 24-hour cultivation. After centrifugation, the cell pellet was collected, and PHB was found to be exclusively accumulated intracellularly. Qualitative analysis of PHB was carried out using FT-IR, confocal laser scanning microscopy (LSCM), transmission electron microscopy (TEM), and GC–MS. Quantification was performed by GC–MS. Detailed procedures for cultivation and analysis are provided in the Supplementary Materials 14 to 18.

### Genome and Transcriptome Analysis for LETBE-HOU

Samples for genomic DNA and transcriptomic RNA extraction were prepared by culturing cells in LB and YSV media for 24 h, respectively, followed by centrifugation and immediate flash-freezing. For transcriptomic, the control group was not supplemented with PE degradation products, whereas the experimental group received 1 g/L of these substrates. Both DNA and RNA extraction, as well as quality control, were conducted by Guangzhou GediO.

The genomic data were functionally annotated through GO, KEGG, COG, PHI, and CAZy databases. The complete genome sequence has been deposited in the National Center for Biotechnology Information (NCBI) database under accession number CP176001.

For transcriptome analysis, gene expression levels were quantified using RSEM (v1.2.19)^37^. Differentially expressed genes were identified and subjected to KEGG pathway annotation and GSEA to elucidate the PHB synthesis pathway and identify key genes involved.

## Acknowledgements

This work was supported by the National Key Research and Development Program of China (2019YFA0706900), and Jiangsu Provincial Science and Technology Department Policy Guidance Program-International Cooperation Projects-Innovation cooperation project of “B&R” (No. BZ2020010).

## Author contributions

J.W. initiated the project. D.M.K., L.W., W.X., M.S., and G.Y.Z., performed all the experiments, D.M.K. performed the computational work, D.M.K. drafted the manuscript, which was revised and approved by all authors.

