## Supplemental Information for "Upcycling Polyethylene into Poly(3-hydroxybutyrate) via a Chemo-Enzymatic-Microbial Cascade"

### Supplementary Methods

#### 1. Reagents, strain, and plasmid

Polyethylene (PE) was obtained from Macklin. Most Chemicals were sourced from TCI, Sinopharm or Macklin. Plasmid construction was performed using *Escherichia coli* (*E. coli*) JM109 as the cloning host, while heterologous protein expression was carried in *E. coli* BL21(DE3), *B. paranthracis* LETBE-HOU was obtained through the screening of wild strain. The recombinant plasmid pET-24a(+)-*tfcut*, encoding the cutinase from *Thermobifida fusca* (*Tf*Cut) or its mutants, was either preserved from previous work[1] or constructed therein. The *Tf*Cut protein used in this study was engineered in our laboratory by introducing the mutations: D204C/E253C/F209I/Q92G/H184S/I213K into the wild-type sequence[1]. Additionally, the plasmid pUB110-phaC was constructed in this study to encode the 3-hydroxybutyrate polymerase (PhaC) gene from the wild-type strain *B. paranthracis* LETBE-HOU.

#### 2. Expression, mutation,and purification of *Tf*Cut

*E. coli* cultures were grown in LB medium at 37°C with shaking at 200 rpm for 10 hours. Fermentation was subsequently carried in TB medium (containing 30 μg/ mL kanamycin). After 2 hours of incubation at 37°C and 200 rpm, the temperature was lowered to 25°C.Protein expression was induced by adding 0.1 mM IPTG, and cultures were incubated for an additional 24 hours. Cells were then harvested by centrifugation, and the supernatant containing the secreted target protein was collected.

Site-directed mutagenesis was performed via PCR using pET-24a(+)-*tfcut* plasmid as the template and the primers listed in Table S1. For constructing iterative combinatorial mutants, the plasmid encoding the most promising variant from the previous screening round served as the template for the next round. All *Tf*Cut and its variants contained a C-terminal His-tag for protein purification. Protein purification was performed using Ni-NTA affinity chromatography. The purification buffers employed were: Buffer A (25 mM Tris-HCl buffer, 500 mM NaCl, pH 7.4) and Buffer B (25 mM Tris-HCl buffer, 500 mM NaCl, 500 mM imidazole, pH 7.4). After applying the protein sample to the Ni-NTA resin, the column was washed with 10 column volumes of 100% Buffer A, followed by step-wise elution with buffers containing 5% and 20% Buffer B (in Buffer A). The final eluate was collected, and imidazole was removed via ultrafiltration and buffer exchange. The resulting purified protein was used for subsequent enzymatic characterization.

##### 3. Characterization of *Tf*Cut

The optimal temperature of *Tf*Cut and its mutants was determined across a range of 50°C to 95°C in 5°C increments. Relative activity at each temperature was calculated as a percentage of the maximum observed activity. Similarly, the optimal pH was assessed using buffers spanning pH 6.0 to 10.0 in 0.5-unit intervals, with relative activity expressed relative to the peak value. The thermal half-life ($t_{1/2}$) was determined by incubating the enzyme at 70°C and pH 9.5, and calculated based on first-order inactivation kinetics according to the method of Rao et al[2]. The melting temperature ($T_m$) was measured by differential scanning microcalorimetry using a protein concentration of 1.5 mg/mL. Scans were performed from 30°C to 110°C at a heating rate of 1°C/min under 3 atm pressure.

##### 4. Pre-treatment of PE

The pre-treatment procedure was as follows: (1) A mixture of equal volumes of 1,2,4-trichlorobenzene and 1,2-dichloroethane was prepared, to which 5% (w/v) PE was added as substrate. The mixture was stirred and dissolved at 100°C and 400 rpm for 12 h. (2) The catalyst $^{m}$CPBA was added at a 1:5 mass ratio (catalyst to PE), and the reaction proceeded at 90°C with stirring at 400 rpm for 8 h; (3) Two volumes of methanol were added to precipitate the pre-treated PE, which was then collected by filtration. This precipitation and washing step were repeated 3–4 times to remove residual solvents and catalyst. The sample was air-dried in a fume hood to volatilize residual organic reagents, followed by drying in a 60°C oven until a constant weight was achieved.

##### 5. Degradation of pre-treated PE

Pre-treated PE was added at 1% (w/v) to the *Tf*Cut enzyme solution. The reaction was incubated at 60°C with shaking at 200 rpm in a water bath for 7 days. After the reaction, the mixture was collected and subjected to solid-liquid separation. The solid product was sequentially washed via ultrasonication for 30 min with 2% (w/v) SDS, 75% (v/v) ethanol, and deionized water. The product was then dried in a 60°C oven until constant weight.

During the optimization of degradation parameters, the following key factors were investigated: additive type and concentration, enzyme loading, reaction temperature, pH, and duration. The specific optimization conditions included: (1) Additive type: anionic surfactant (SDS), cationic surfactant (CTAB), nonionic surfactant (Triton X-100, PEG400), biosurfactant (betaine or rhamnolipid), encapsulant (α-CD, β-CD, or γ-CD), and organic solvent (DMSO); (2) Additive concentration (g/L): 0.12, 0.24, 0.36, 0.48, 0.60, 1.20, 2.40, and 3.60; (3) Enzyme loading (U/mg substrate): 1, 2, 3, 4, 5, 10, 20, 30, and 40; (4) Reaction temperature (°C): 50, 55, 60, 65, 70, 75, and 80; (5) Reaction solution pH: 7.0, 7.5, 8.0, 8.5, 9.0, 9.5, and 10.0; (6) Reaction duration (days): 1, 2, 3, 4, 5, 6, and 7.

#### 6. Random Forest Modeling and Analysis

The random forest model was constructed using the randomForest package in R. The independent variables comprised additive type and concentration, enzyme loading, reaction temperature, pH, reaction time, rotational speed, and the ester bonds ratio of pretreated polyethylene, while weight loss rate served as the dependent variable. To mitigate scaling effects among predictors, temperature, enzyme loading, and rotational speed were log10-transformed prior to modeling. Two key hyperparameters, the number of trees (ntree) and the number of predictors sampled at each split (mtry), were systematically optimized. The ntree parameter was evaluated across 25 gradients from 200 to 5000 trees (in increments of 200), and mtry was tested across 8 gradients from 1 to 8. A total of 200 parameter combinations were screened, with each combination simulated 10 times. During each simulation, 90% of the data were randomly selected as the training set and the remaining 10% were used as the test set to evaluate model accuracy. The optimal ntree and mtry values were determined based on the results from 10 replicate simulations.

For model interpretation, the vivid[3] package was employed to analyze variable importance (Vimp), variable interaction measures (Vint), partial dependence plots

(PDP), and individual conditional expectation (ICE) curves. SHapley Additive exPlanations analyses (SHAP) analysis was conducted using the shaply[4] package to quantify the contribution of each independent variable to model predictions.

**7. Weight loss rate analysis of PE degradation products**

The reaction solution was filtered using a 600-mesh nylon mesh. The retained solid particles were washed, dried, and weighed. The weight loss ratio ($\Delta W$) was calculated using the following Formula 1. $\Delta W$ was weight loss ratio, $S$ was the weight of nylon mesh and products, $S_0$ was the weight of nylon mesh, $M$ was the weight of substrate, $s_0$ and $s_1$ was the weight of empty nylon mesh before and after drying. The enzyme activity residual rate ($\Delta E$) was calculated using Equation 2, where $E_0$ and $E_1$ represent the initial enzyme activity and the enzyme activity after the reaction, respectively.

$$\Delta W=\frac{S\text{-}S_0}{M}\times\frac{s_0\text{-}s_1}{s_0} \quad \text{Equation 1}$$

$$\Delta E=\frac{E_1}{E_0} \quad \text{Equation 2}$$

**8. Differential Scanning Calorimeter (DSC)**

Thermal analysis was performed using differential scanning calorimetry under a nitrogen atmosphere. Samples were heated from 30°C to 300°C at a rate of 10°C/min. The melting temperatures of PE under different catalytic conditions were determined using TA Instruments Universal Analysis 2000 software.

**9. Fourier Transform Infrared Spectroscopy (FT-IR)**

Both *Tf*Cut degradation products and LETBE-HOU metabolites were analyzed using the same FT-IR method. Fourier transform infrared spectra were acquired in transmission mode with 32 scans over the spectral range of 4000–600 $cm^{-1}$ at a resolution better than 0.5 $cm^{-1}$ (signal-to-noise ratio > 44000:1). Data were processed using Omnic 8.2.0.387 software, with baseline correction and peak area analysis performed in absorbance mode.

**10. High-Temperature Gel Permeation Chromatography (HT-GPC)**

Molecular weight distributions were determined by high-temperature gel permeation chromatography. Polyethylene samples were dissolved in 1,2,4-trichlorobenzene at a final concentration of 2 mg/mL. Separation was achieved using TSKgel GMH H(S)

HT2 and TSKgel H HFRC tandem columns equipped with a TSK guardcolumn H(S) HT2 guard column. The analysis was conducted at 145°C with an injection volume of 300 μL, using 1,2,4-trichlorobenzene as the mobile phase at a flow rate of 1.0 $mL \cdot min^{-1}$ over a 50-minute elution time. Data were processed by min-max normalization and presented as differential distribution curves (dW/d(log M) versus log M).

**11. Gas chromatography–mass spectrometry (GC–MS)**

Volatile products from the enzymatic degradation of PE by *Tf*Cut were analyzed using an Agilent GC–MS system equipped with a DB-5MS column (30 m × 0.25 mm × 0.25 μm). The GC was operated under the following conditions: injector temperature, 250°C; injection volume, 1 μL; helium carrier gas at a flow rate of 1 mL/min; split ratio, 10:1. The oven temperature was programmed from 40°C (no hold) to 250°C at a rate of 10°C/min, followed by a 4-minute hold. Mass spectrometric detection was performed with a mass range of 33–450 m/z, an ion source temperature of 250°C, and an electron energy of 70 eV.

**13. Wild bacteria screening process**

(1) Sample Collection

Seven environmental samples were collected for wild strain screening. These included adhesive deposits from a paper mill in Guangrao County, Shandong Province (Samples 1 and 2), and soil samples from plastic-contaminated agricultural regions across China: Binhu District, Wuxi City, Jiangsu Province (Samples 3 and 4); Jinzhou City, Shijiazhuang, Hebei Province (Sample 5); Gaotang County, Liaocheng City, Shandong Province (Sample 6); and Heishi Town, Panshi City, Jilin Province (Sample 7). Samples 1 and 2 contained ester bond-rich adhesives, while Samples 3–7 were obtained from farmland with historical PE mulch film pollution.

(2) Enrichment Culture in Liquid Medium

Each sample was introduced at 2 g/L into an enrichment medium (NaCl 2.0 g/L, $K_2HPO_4$ 0.7 g/L, $KH_2PO_4$ 0.7 g/L, $MgSO_4 \cdot 7H_2O$ 0.7 g/L, $NH_4SO_4$ 1.0 g/L, $FeSO_4 \cdot 7H_2O$ 0.002 g/L, $ZnSO_4 \cdot 7H_2O$ 0.002 g/L, and $MnSO_4 \cdot 5H_2O$ 0.001 g/L) as the microbial inoculum, supplemented with 1 g/L PE degradation products as the sole carbon source. Cultures were incubated at 37°C with shaking at 200 rpm for 10 days.

(3) Liquid Medium Subculturing

Following the primary enrichment, 2 mL of culture was aseptically transferred into fresh enrichment medium containing 1 g/L PE degradation products. The subculture was incubated under identical conditions (37°C, 200 rpm) for an additional 10 days.

(4) Isolation on Solid Medium (Primary Screening)

A solid screening medium (adding 1.8% agar powder to the enrichment medium) was prepared by adding 1 g/L PE degradation products and 1 mg/L Nile red to the base agar. After subculturing, 1 mL of culture was centrifuged (3,000 rpm, 5 min). The pellet was resuspended, serially diluted in enrichment medium, and spread onto the solid medium. Plates were incubated inverted at 37°C in the dark for 7 days. All plating procedures were conducted under subdued light to minimize photobleaching of Nile red.

(5) Metabolic Capability Assay (Secondary Screening)

Individual colonies were inoculated into LB medium and grown at 37°C with shaking at 200 rpm for 12 h. A 5% (v/v) inoculum was then transferred to YSV medium[5] containing 1 g/L PE degradation products and cultivated at 30°C with shaking at 200 rpm for 24 h. After cultivation, residual PE particles were removed by filtration through a 30 μm hydrophobic membrane. Bacterial cells were harvested from the filtrate by centrifugation at 8,000 rpm, and intracellular PHA accumulation was quantified via Nile red staining[6] and fluorescence measurement (550 nm excitation and 590 nm emission)[7].

**14. FT-IR qualitative of poly-3-hydroxybutyrate (PHB)**

For FT-IR analysis, PHB was first extracted from the cellular biomass using a previously described protocol[8]. The extracted polymer was then analyzed by FT-IR, following the same instrumental method as detailed elsewhere in this study, to confirm its chemical identity.

**15. Laser scanning confocal microscopy (LSCM) qualitative of PHB**

Following three washes with physiological saline, the cells were stained with Nile red and visualized by LSCM. Images were acquired at 7,000× total magnification under oil immersion, using 540 nm excitation and 590 nm emission wavelengths to detect the green fluorescence[7].

**16. Transmission Electron Microscope (TEM) qualitative of PHB**

Following centrifugation (12,000 rpm, 1 min) of 10 mL culture, the cell pellet was washed thrice with 0.9% (w/v) saline and fixed in 2.5% (v/v) glutaraldehyde at 4°C. Cell embedding and transmission electron microscopy were subsequently carried out by Beijing Yangou, with imaging at 3,000× and 30,000× magnification.

**17. GC–MS qualitative and quantification of PHB**

The PHB content in LETBE-HOU was quantified via GC–MS following a derivatization procedure adapted from Liu et al[9]. The polymeric PHB was first hydrolyzed and methanolized to convert 3-hydroxybutyrate monomers into volatile methyl esters.

A six-point calibration curve was established using purified PHB as an external standard (ESTD) at concentrations of 12, 32, 70, 113, 141, and 199 mg/L, with methyl benzoate (117 mg/L) as the internal standard (ISTD). The calibration curve was generated by plotting the ratio of the ESTD-to-ISTD peak areas against the ratio of the ESTD-to-ISTD concentrations (Fig. S14). The PHB concentration in experimental samples was determined by interpolating the measured peak area ratio into this calibration curve.

For sample pretreatment, lyophilized cell pellets (about 20 mg) were transferred into esterification tubes. Then, 2 mL of 15% (v/v) sulfuric acid-methanol solution and 2 mL of a 200 mg/L methyl benzoate in chloroform were added. The tubes were heated at 105°C for 3 h, then cooled on ice for 10 min. Thereafter, 1 mL of deionized water was added for back-extraction. After vortexing for 10 min and centrifugation, the organic phase was collected, diluted with chloroform to an appropriate concentration, and dehydrated over anhydrous sodium sulfate. The extract was finally filtered through a 0.22 μm organic solvent-resistant membrane prior to GC–MS analysis.

GC–MS analysis was performed using a TG-5SILMS column (30 m × 0.25 mm × 0.25 μm). The injector temperature was set to 250°C, with an injection volume of 1 μL and a split ratio of 10:1. High-purity helium was used as the carrier gas at a constant flow rate of 1.0 mL/min. The oven temperature program was as follows: initial temperature 50 °C, increased to 110°C at a rate of 10°C/min, then raised to 230°C at 13.3°C·min$^{-1}$ and held for 6 min. The mass spectrometer was operated in electron ionization mode at 70 eV, with an ion source temperature of 230 °C and a mass scan range of 33–350 m/z.

**18. Overexpression of the LETBE-HOU gene**

The overexpression plasmid pUB110-phaC, carrying the phaC gene from Bacillus paranthracis LETBE-HOU, was constructed using overlap extension PCR. The *phaC* gene was amplified from the genomic DNA of *B. paranthracis* LETBE-HOU using primers phaC-F/phaC-R (shown in Table S1), while the pUB110 backbone was amplified from the commercial plasmid using primers pUB-F/pUB-R. These two fragments were then assembled by overlap extension PCR. The resulting recombinant plasmid was first cloned in B. subtilis SCK6. After verification by Sanger sequencing, the correct plasmid was subsequently transformed into the wild-type strain B. paranthracis LETBE-HOU.

### Supplementary Figs. 1-16

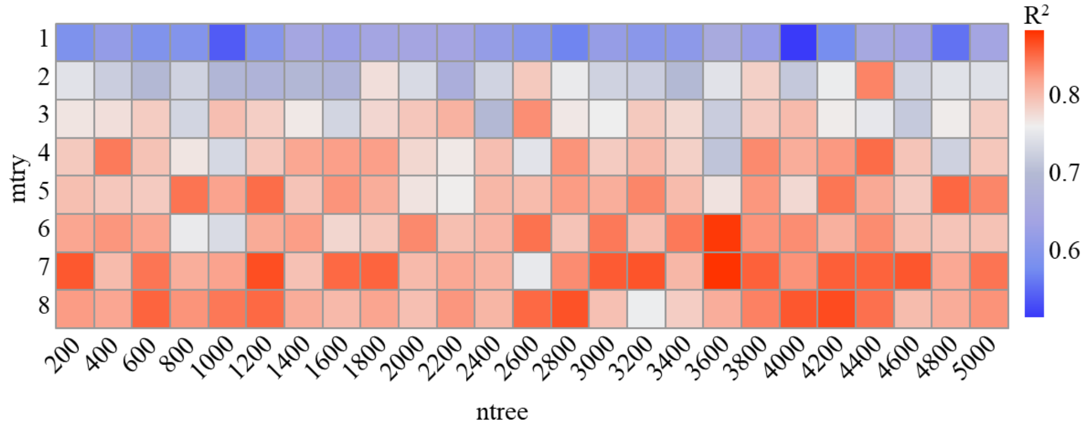


**Supplementary Fig. 1. Parameter screening for random forest model.**

The $R^2$ value represents the Pearson correlation coefficient, indicating the agreement between actual and predicted values across ten modeling runs for each mtry and ntree parameter combination. A blue-white-red color gradient is used to visualize $R^2$ values, with blue, white, and red corresponding to low, intermediate, and high values, respectively.

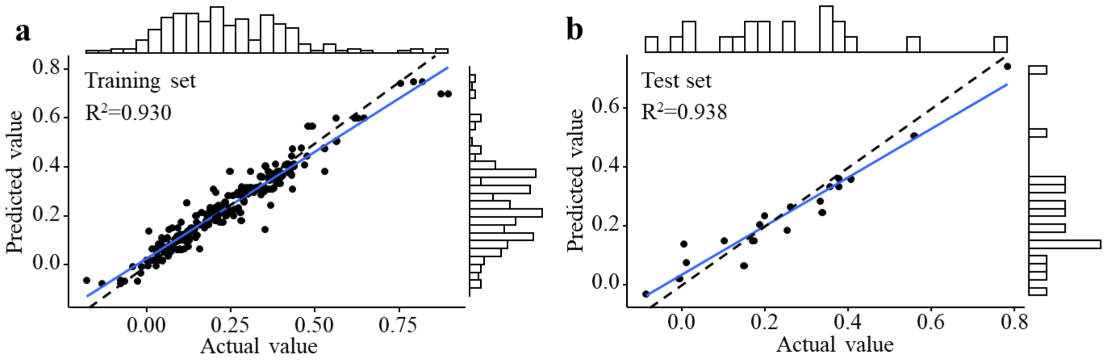


**Supplementary Fig. 2. Visualization analysis of regression performance for the random forest model.**

**a,** Training set performance. **b,** Test set performance. Frequency histograms represent the distributions of observed versus predicted values.

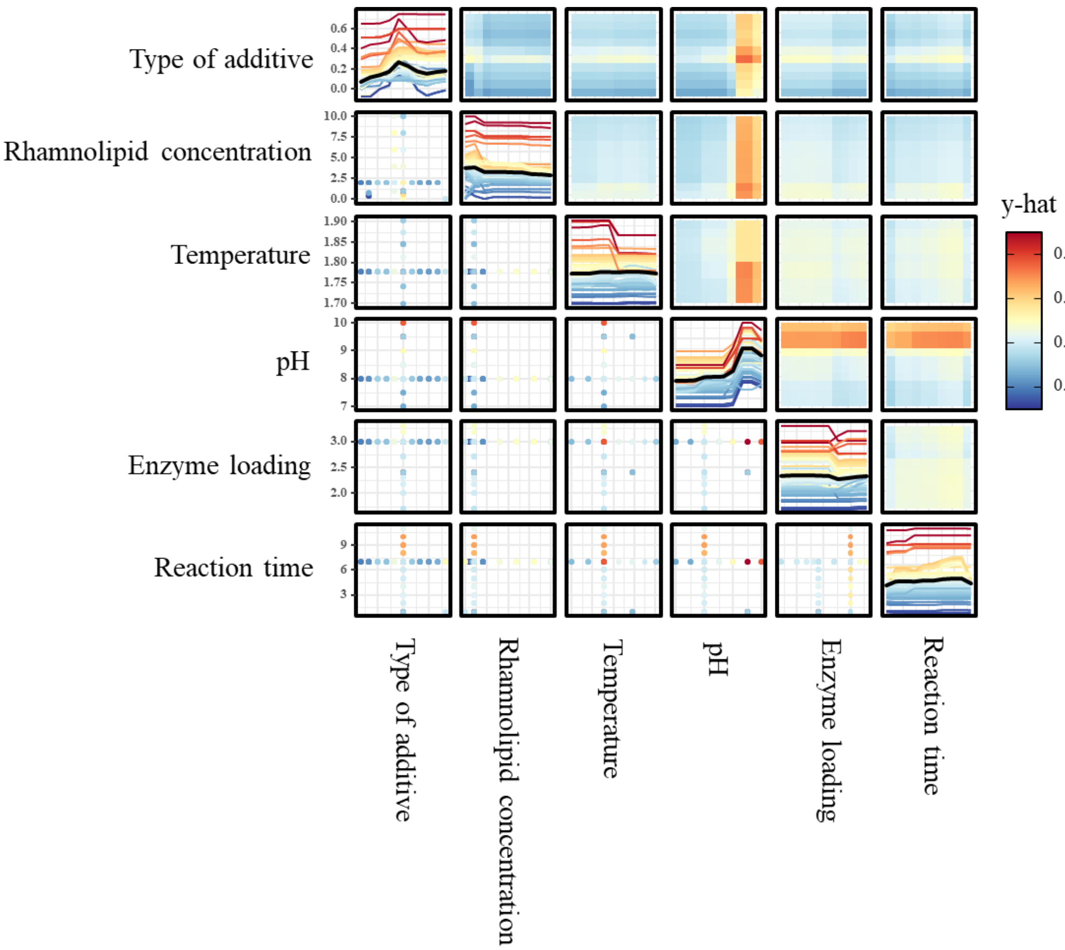


**Supplementary Fig. 3. Vivid package analysis results of the random forest model.** Colored lines along the diagonal represent ICE analysis, while the overlaid black lines indicate PDP. The upper triangular panels show bivariate PDP, and the lower triangular panels display data distribution scatter plots.

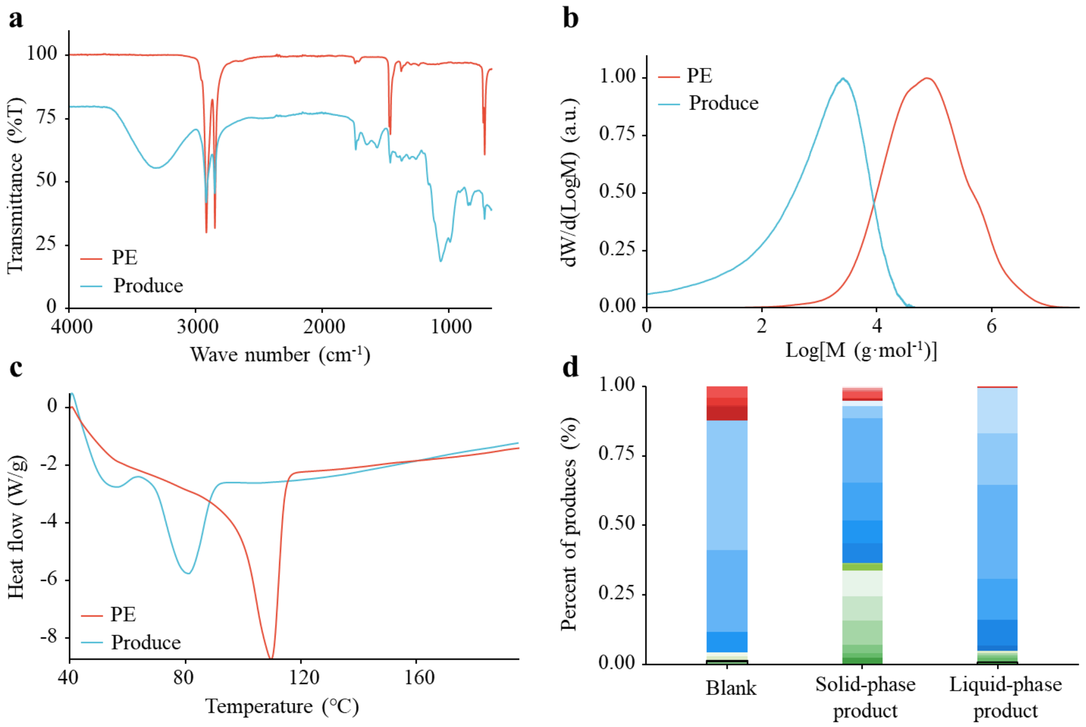


**Supplementary Fig. 4. Product characterization of *Tf*Cut degradation system under optimized conditions.**

**a,** FT-IR spectrum showing introduced oxygen-containing functional groups. **b,** HT-GPC profile revealing molecular weight distribution. **c,** DSC thermogram indicating thermal behavior. **d,** GC–MS chromatogram with chemical classes color-coded as follows: monocarboxylic acids (red), dicarboxylic/hydroxy fatty acids (yellow), diols (blue), and monoalcohols (green).

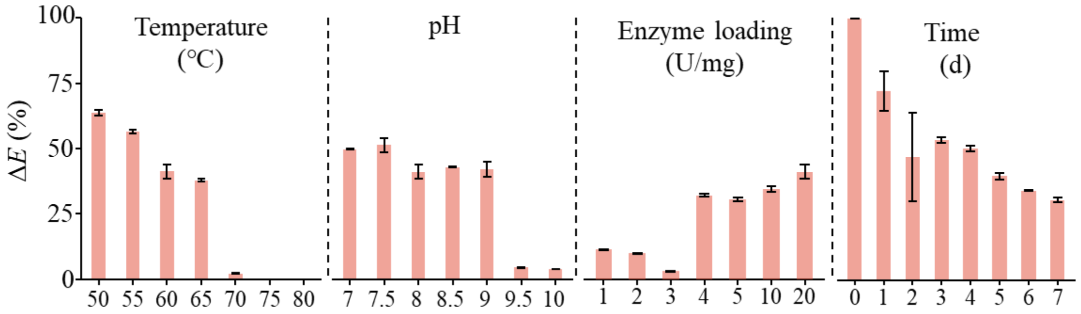


**Supplementary Fig. 5. Enzyme activity residual rate of *Tf*Cut across different reaction conditions.**

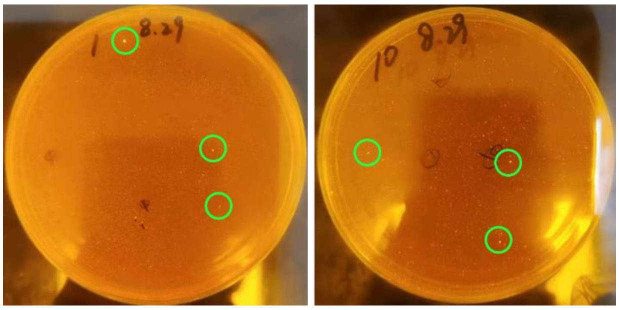


**Supplementary Fig. 6 Primary screening of wild-type strains on solid medium with polyethylene degradation intermediates as the sole carbon source.**

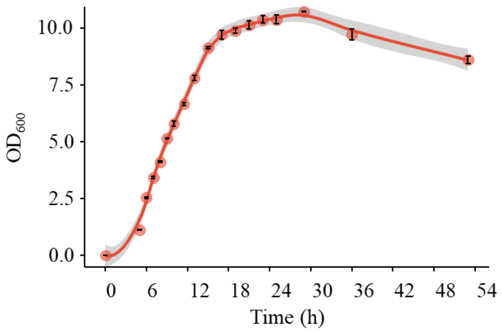


**Supplementary Fig. 7 Growth curve of LETBE-HOU in LB medium at 37°C.**

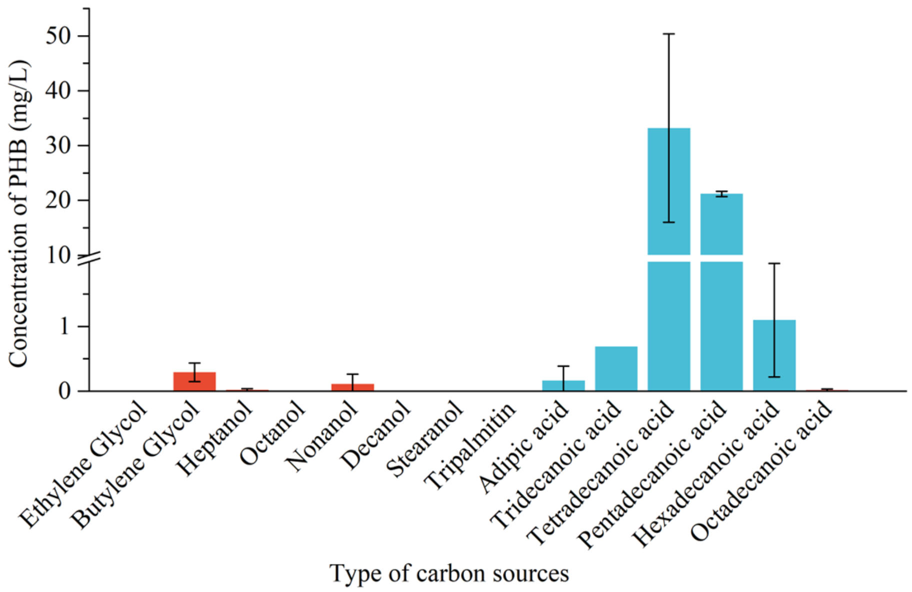


**Supplementary Fig. 8 Carbon source utilization profile of LETBE-HOU.**

Red bars: alcohol substrates; blue bars: acid substrates.

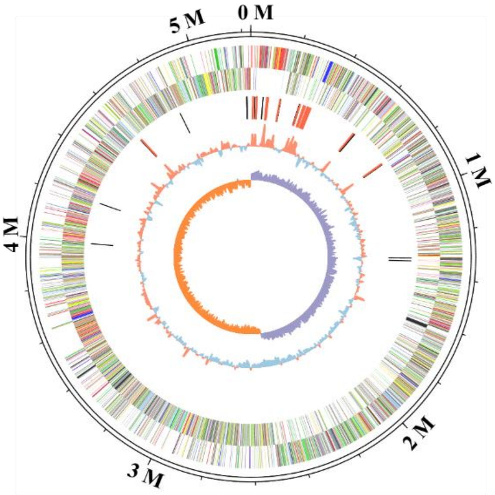


**Supplementary Fig. 9 Circular genome map of LETBE-HOU.**

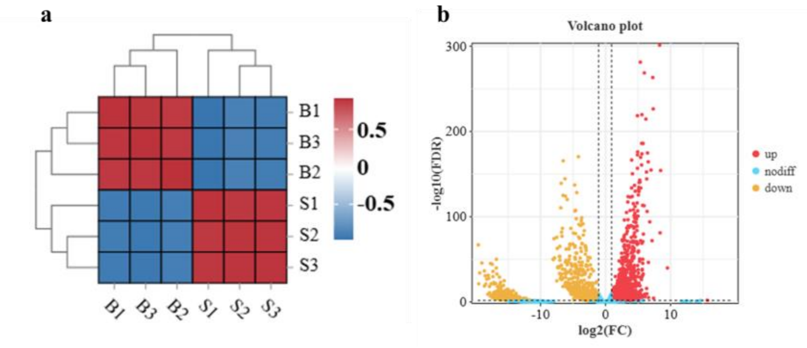


**Supplementary Fig. 10 Comparative transcriptomic analysis of LETBE-HOU**

**a,** Inter-sample correlation analysis. **b,** Differential gene expression between control (B) and experimental (S) groups.

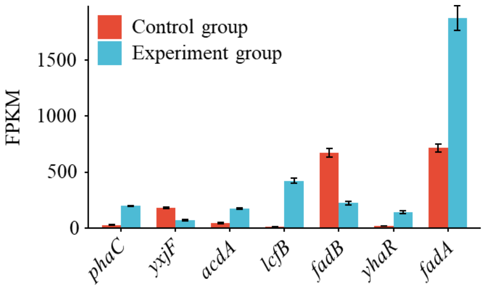


**Supplementary Fig. 11 Expression level of key gene in PHB biosynthesis pathway by LETBE-HOU**

**Table S1 Genome overview of LETBE-HOU**

| Features | Chromosome |
|---|---|
| Size (bp) | 5,234,090 |
| GC content (%) | 35.55 |
| Protein coding genes (CDSs) | 5198 |
| 5S_rRNA | 14 |
| 23S_rRNA | 14 |
| 16S_rRNA | 14 |
| tRNAs | 105 |
| sRNAs | 12 |
